# Loss of function mutations in *epaR* confer resistance to phage NPV1 infection in *Enterococcus faecalis* OG1RF

**DOI:** 10.1101/301929

**Authors:** Khang Ho, Wenwen Huo, Savannah Pas, Ryan Dao, Kelli L. Palmer

**Affiliations:** Department of Biological Sciences, The University of Texas at Dallas

## Abstract

*Enterococcus faecalis* is a Gram-positive opportunistic pathogen that inhabits the human gastrointestinal tract. Because of the high frequency of antibiotic resistance among *Enterococcus* clinical isolates, interest in using phage to treat enterococcal infections and to decolonize high-risk patients for antibiotic-resistant *Enterococcus* is rising. Bacteria can evolve phage resistance, but there is little published information on these mechanisms in *E. faecalis*. In this report, we identified genetic determinants of *E. faecalis* resistance to ϕNPV1. We found that loss-of-function mutations in *epaR* confer ϕNPV1 resistance by blocking phage adsorption. We attribute the inability of the phage to adsorb to the modification or loss of an extracellular polymer in strains with inactivated *epaR*. Phage-resistant *epaR* mutants exhibited increased daptomycin and osmotic stress susceptibilities. Our results demonstrate that *in vitro* spontaneous resistance to ϕNPV1 comes at a cost in *E. faecalis* OG1RF.

## Introduction

*Enterococcus faecalis* is a Gram-positive bacterium that inhabits the human gastrointestinal tract and is associated with nosocomial infections (1). Infections caused by *E. faecalis* can be difficult to treat because of the high frequency of resistance to multiple antibiotics among *E. faecalis* clinical isolates (2). The antibiotic daptomycin can be used to treat certain infections caused by multidrug-resistant enterococci. Daptomycin is a lipopeptide antibiotic that interacts with the enterococcal cell surface and disrupts membrane structure and function (3).

Bacteriophages (phages) are bacterial viruses and natural predators of bacteria. It is reasonable to expect that phages can be employed to treat bacterial infections. However, phages have not been extensively studied in the Western world in the context of therapeutic application until recently due to the availability of antibiotics (4). In recent years, interest in using phages to treat bacterial infections (phage therapy) has reemerged because of the emergence of multidrug-resistant bacteria. For *E. faecalis*, promising studies include the use of phage to eliminate biofilm, a major barrier to antibiotic treatment, and to increase survival rates in mouse models of enterococcal infection (5, 6).

One advantage of phage therapy is limited damage to the native microbiome because of the specificity of the phage to its host (7). Typically, lytic phage have narrow host ranges and are species-specific or target a range of strains within a species. The first step to a successful phage infection is the attachment of the phage particle to the proper receptor present on the surface of the host cell. Phage receptors have been extensively studied in certain phage families, including the T series phages, Mu, and λ for Gram-negative bacteria (8-11). Some phage receptors have been characterized in Gram-positive bacteria, including receptors for ϕSPP-1 of *Bacillus subtilis* (12) and the phage c2 group of *Lactococcus lactis* (13, 14). YueB, the ϕSPP-1 receptor, and PIP, the phage c2 receptor, are orthologs and are required for irreversible phage adsorption (12). Enterococcal phage receptors have not been well-characterized. Previously, we and collaborators identified PIP as a receptor and potential DNA channel for the *E. faecalis* phages ϕVPE25 and ϕVFW (15).

Bacteria can evolve phage resistance. Mechanisms of phage resistance include modification or loss of the phage receptor (16). However, as phage receptors generally serve physiological functions in the cell, the modification or loss of a receptor could come at a cost for the bacterial host. For example, spontaneous phage-resistant mutants have altered antibiotic sensitivity in *P. aeruginosa* (17). Phages utilizing receptors that have roles in antibiotic resistance could be advantageous for re-sensitizing resistant bacteria to antibiotics.

Considering the increasingly limited treatment options for *E. faecalis* infections and the revival of interest in using phage therapy to treat bacterial infections, it is crucial that we know the receptor(s) of enterococcal phages since effective phage cocktails use phages targeting multiple different receptors (18). Moreover, the roles of these receptors in enterococcal physiology should be elucidated. The tailed, virulent phage ϕNPV1 was found in previous studies to infect *E. faecalis* OG1RF (19, 20). In this study, we used a combination of genomic and genetic approaches to investigate the ϕNPV1 receptor in *E. faecalis* OG1RF.

## Results

### Deletion of *epaR* alters susceptibility to ϕNPV1

We isolated a OG1RFΔPIP (15) strain with spontaneous resistance to ϕNPV1 (Figure 1). We refer to this strain as OG1RF-C. The genome sequence of OG1RF-C was determined. We identified non-synonymous substitutions in *epaR, bgsB, iolA2*, and OG1RF_10252 (Table 2). *epaR* is one of the 18 conserved genes of the *epa* gene cluster (*epaA-epaR*), which codes for synthesis of the enterococcal polysaccharide antigen (Epa) (20). *epaR* encodes a putative glycosyltransferase with 5 predicted transmembrane domains, and its role in Epa biosynthesis has not been investigated. The product of *bgsB* is a putative cytoplasmic protein catalyzing the transfer of glucose from UDP-glucose to diacylglycerol (DAG) to form monoglucosyl-DAG. An additional glucose is added to glucosyl-DAG by *bgsA*, forming diglucosyl-DAG. From diglucosyl-DAG, the polymerization of glycerol-phosphate can occur, resulting in lipoteichoic acid (LTA) (21). *iolA2* is predicted to encode a methylmalonate-semialdehyde dehydrogenase, which catalyzes the breakdown of malonic semialdehyde to acetyl-CoA and CO_2_ (22). OG1RF_10252 is predicted to encode a Acyl-ACP_TE domain (pfam01643; e-value 7.510e^−114^) which catalyzes the termination of fatty acyl group extension by hydrolyzing an acyl group on the fatty acid.

**Figure 1.**
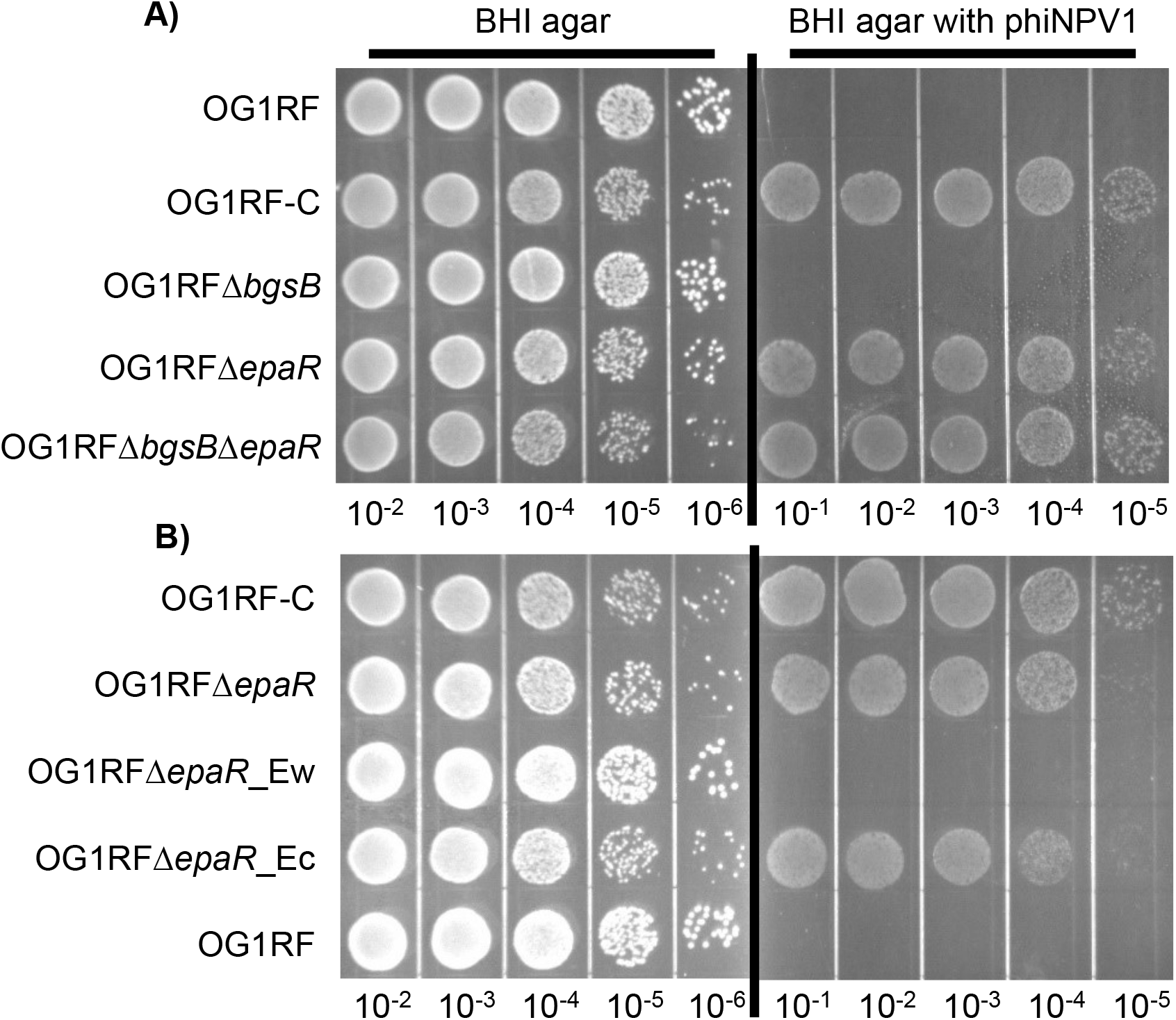
Phage susceptibility assays. Overnight cultures were diluted in PBS and spotted on BHI plates with or without 10^9^ PFU/mL phiNPV1. Images were taken after 18 h incubation at 37°C. The image shown is representative of three independent trials. (A) phiNPV1 susceptibility of *E. faecalis* OG1RF and derivatives. (B) phiNPV1 susceptibility of complemented strains of OG1RF *ΔepaR*.

**Table 1.** Strains and plasmid used.

| Strains and plasmids | Description | Reference |
| --- | --- | --- |
| ***E. faecalis* strains** |  |  |
| OG1RF | Human oral cavity isolate | (40, 41) |
| OG1RFΔPIP | PIP deletion strain | (15) |
| OG1RF-C | OG1RFΔPIP ϕNPV1-resistant strain | This work |
| OG1RFΔ*bgsB* | OG1RF *bgsB* deletion mutant | This work |
| OG1RFΔ*epaR* | OG1RF *epaR* deletion mutant | This work |
| OG1RFΔ*bgsB*Δ*epaR* | OG1RF *bgsB* and *epaR* double deletion mutant | This work |
| OG1RFΔ*epaR*_Ew | OG1RFΔ*epaR* with wild-type *epaR* complementation *in cis* | This work |
| OG1RFΔ*epaR*_Ec | OG1RFΔ*epaR* complemented with OG1RF-C *epaR* allele *in cis* | This work |
| OG1RFΔ*bgsB*_Bw | OG1RFΔ*bgsB* with wild-type *bgsB* complementation *in cis* | This work |
| OG1RFΔ*bgsB*_Bc | OG1RFΔ*bgsB* complemented with OG1RF-C *bgsB* allele *in cis* | This work |
| OG1RF_R2 | OG1RF ϕNPV1-resistant strain | This work |
| OG1RF_R3 | OG1RF ϕNPV1-resistant strain | This work |
| OG1RF_R4 | OG1RF ϕNPV1-resistant strain | This work |
| OG1RF_R5 | OG1RF ϕNPV1-resistant strain | This work |
| OG1RF_R6 | OG1RF ϕNPV1-resistant strain | This work |
| OG1RF_R7 | OG1RF ϕNPV1-resistant strain | This work |
| OG1RF_R8 | OG1RF ϕNPV1-resistant strain | This work |
| OG1RF_R9 | OG1RF ϕNPV1-resistant strain | This work |
| OG1RF_R10 | OG1RF ϕNPV1-resistant strain | This work |
| ***E. coli* strains** |  |  |
| EC1000 | Cloning host. Provides *repA* *in trans* | (42) |
| **Plasmids** |  |  |
| pLT06 | Cloning vector, temperature sensitive *repA*, Cam<sup>r</sup> | (36) |
| pLT06_Δ*bgsB* | *bgsB* deletion construct | This work |
| pLT06_Δ*epaR* | *epaR* deletion construct | This work |
| pLT06_Ew | *epaR* wild-type allele complementation construct | This work |
| pLT06_Ec | *epaR* OG1RF-C allele complementation construct | This work |
| pLT06_Bw | *bgsB* wild-type allele complementation construct | This work |
| pLT06_Bc | *bgsB* OG1RF-C allele complementation construct | This work |

**Table 2.** SNPs detected in OG1RF-C strain.

| Position | Base pair change | Amino acid change | Annotation |
| --- | --- | --- | --- |
| 1800903 | C→T | D361N | *epaR* |
| 1878245 | G→T | S453Y | *bgsB* |
| 2308650 | C→A | G134* | *iolA* |
| 260665 | Insertion of A | H44fs | OG1RF_10252 |

To begin to elucidate the roles of these genes in ϕNPV1 susceptibility, we constructed in-frame deletions of *epaR* and *bgsB*, generating strains OG1RFΔ*epa*R and OG1RFΔ*bgs*B, respectively, and a double deletion strain, OG1RF*ΔepaRΔbgsB*. Phage susceptibility of each of these mutants was assayed. Deletion of *epaR* alone was sufficient to confer phage resistance (Figure 1). In contrast, deletion of *bgsB* alone did not alter phage susceptibility. These results indicate that variation in *epaR* is the major factor conferring resistance to ϕNPV1 in OG1RF-C. Since we observed that deletion of *epaR* in OG1RF conferred phage resistance to the same extent as that observed for OG1RF-C, we did not investigate the effects of *iolA2* and OG1RF_10252 on phage resistance in this study.

### The OG1RF-C *epaR* allele confers ϕNPV1 resistance

To determine whether *epaR* mutation is the major contributor to ϕNPV1 resistance in OG1RF-C, we generated strain OG1RFΔ*epaR*_Ec, an OG1RFΔ*epaR* strain complemented *in cis* with the *epaR* allele from OG1RF-C. We also generated strain OG1RFΔ*epaR*_Ew, an OG1RFΔ*epaR* strain with a reconstituted wild-type *epaR*. Complementation with the *epaR* allele of OG1RF-C conferred phage resistance to OG1RFΔ*epaR* (Figure 1B). The wild-type *epaR* allele restored phage susceptibility to OG1RFΔ*epaR* (Figure 1B). Because the mutated *epaR* allele from OG1RF-C confers a phage resistance phenotype, as did deletion of *epaR*, we infer that the *epaR* mutation in OG1RF-C confers loss of function.

### Growth rates of *E. faecalis* strains

We determined generation times for wild-type *E. faecalis* OG1RF, OG1RF-C, and the *epaR* and *bgsB* deletion mutants and complemented strains cultured in BHI broth (Table S1). Average generation times range from a minimum of 27.8 minutes for the *bgsB* deletion mutant to a maximum of 42.7 minutes for the *epaR bgsB* double deletion mutant. The generation times for the wild-type and OG1RF-C strains were similar (30.4 versus 33.9 minutes).

### *epaR* is required for phage adsorption

We were next interested in how *epaR* inactivation protects OG1RF from ϕNPV1 infection. After 15 minutes incubation with ϕNPV1, ~95% of the phage adsorbed to wild-type OG1RF (Figure 2). In contrast, under the same conditions, ~1-2% of the phage adsorbed to OG1RF-C or OG1RFΔ*epaR*. Consistent with the observation that *bgsB* deletion alone confers no significant phage resistance, ~90% of the phage adsorbed to OG1RFΔ*bgsB* within the same experimental settings. These data indicate that *epaR* is required for ϕNPV1 adsorption.

**Figure 2.**
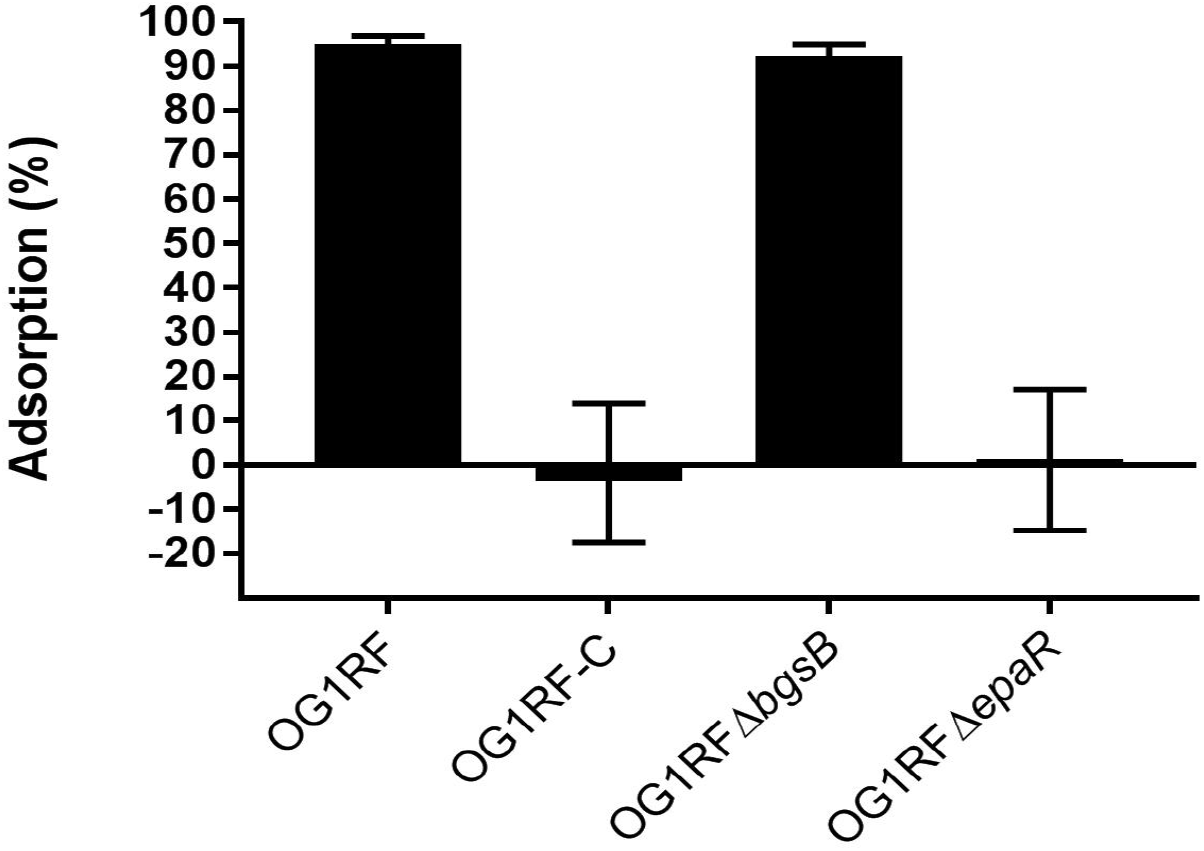
phiNPV1 adsorption assays. Overnight cultures were diluted 1:5 in fresh BHI and equilibrated at 37°C. phiNPV1 was added at a M.O.I. of 10^−2^. After 15 min incubation, 1 mL of each culture was centrifuged, and the supernatant was titered with the phage spot assay. A medium with only phage was used as the control. Percent adsorption was calculated as [(PFU culture – PFU control)/ PFU control] *100%. Data are the average of three independent trials.

### Inactivation of *epaR* alters the Epa polymer

Next, we sought to determine whether the Epa polymer was altered in mutants defective for ϕNPV1 adsorption. The Epa polymer has been extracted and visualized by different groups using different methods (20, 23). We based our method on that from Teng, et al. (20). We found that solubilizing the precipitation with 50% acetic acid improved visualization of the polymers. Gel electrophoresis analysis of carbohydrate extracts found that OG1RF with either an *epaR* deletion or the *epaR* allele from OG1RF-C exhibited loss of a band (P1) that is present in wild-type OG1RF, OG1RFΔ*bgsB*, and the reconstituted *epaR* strain, OG1RFΔ*epaR*_Ew (Figure 3). We conclude that P1 represents an epaR-dependent polymer. Note that the cationic dye Stains-All was used for polymer detection. With the staining methodology used, we cannot state conclusively whether the P1 polymer fails to be synthesized in *epaR* mutants, or if it is synthesized but has a different charge than in wild-type OG1RF. Since NPV1 cannot bind to *epaR* mutants, we hypothesized that the P1 polymer is the phage receptor. However, when we pre-incubated NPV1 with crude carbohydrate extract prior to infection of host cells, we did not observe a decrease in PFU for any extract (Figure S1), suggesting that the polymers in the crude extract are not sufficient for phage adsorption.

**Figure 3.**
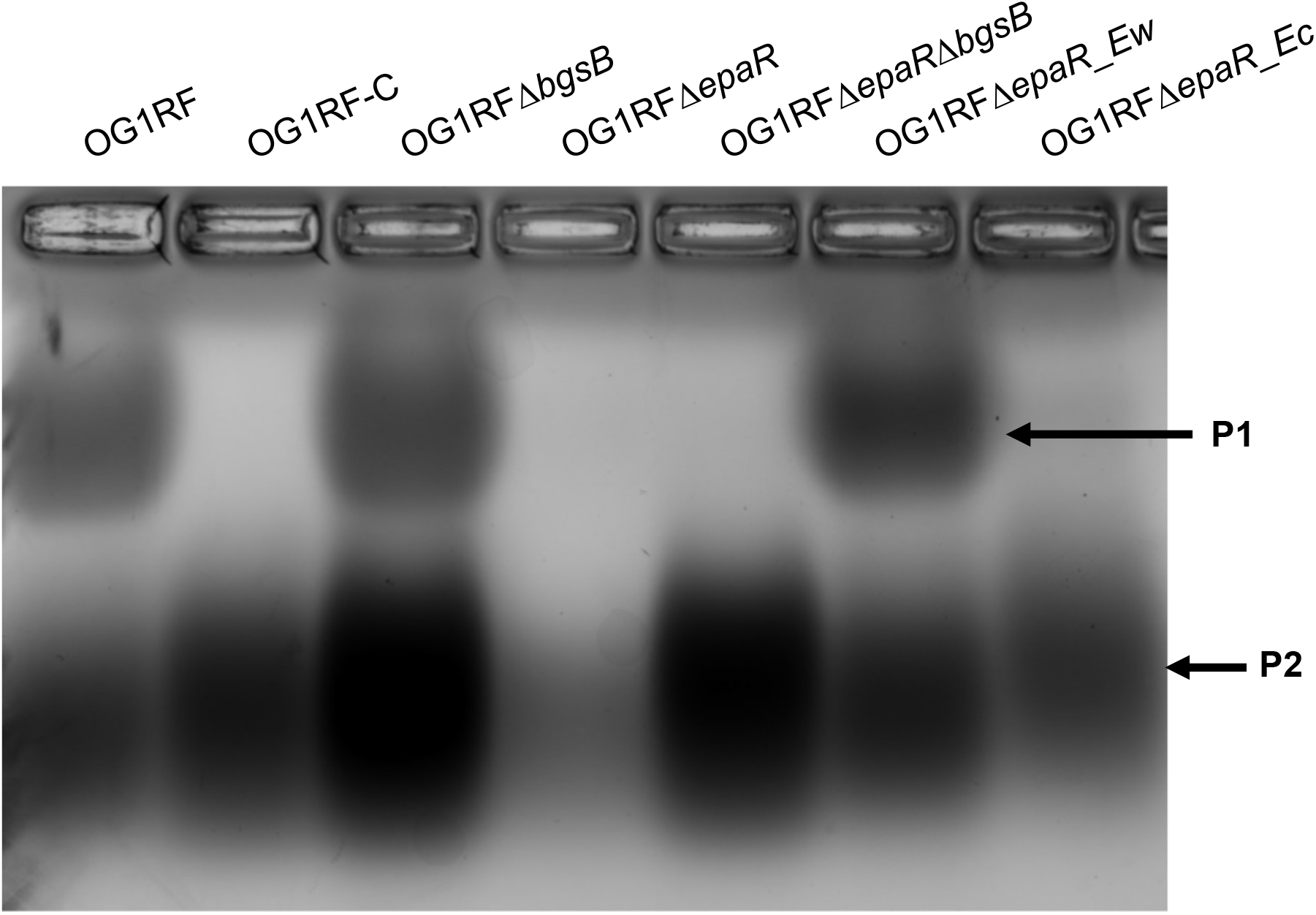
Carbohydrate extract analysis. Carbohydrate was extracted from 200 mL overnight cultures and visualized with Stains-all. The image shown is representative of two independent trials.

We observed an increase in the intensity of the P2 band in OG1RFΔ*bgsB* and OG1RFΔ*bgsB*Δ*epaR* compared to wild-type OG1RF, suggesting that the product P2 is increased in these two deletion strains. Since deletion of *bgsB* in *E. faecalis* results in accumulation of LTA (21), product P2 may represent LTA.

### NPV1-resistant mutants have increased susceptibility to daptomycin

Dale et al. reported increased daptomycin susceptibility in an OG1RF derivative with a deletion in *epaO* (23). Moreover, we identified a *bgsB* mutation in a laboratory-evolved *E. faecium* isolate with decreased daptomycin susceptibility (24). Because of these results, we investigated the daptomycin susceptibilities of our OG1RF mutants (Figure 4). We found that OG1RF-C, OG1RFΔ*epaR*, and OG1RFΔ*epaR* complemented with the OG1RF-C *epaR* allele were each significantly more susceptible to daptomycin than OG1RF. Interestingly, deletion of *bgsB* also conferred increased daptomycin susceptibility (Figure 4). This was complemented by expression of the wild-type *bgsB* allele *in cis*, but not by expression of the OG1RF-C *bgsB* allele *in cis* (Figure 4). Finally, daptomycin susceptibility was substantially altered in the OG1RF*ΔepaR*Δ*bgsB* mutant, with 3 of 6 experimental trials resulting in an MIC below the level of detection of the E-test strip (<0.016 μg/mL; a value of 0.008 μg/mL was used for these data points in statistical analysis). Without complete data regarding the MIC of OG1RF Δ*epaR*Δ*bgsB*, we did not quantitatively determine whether there is a synergistic relationship between *bgsB* and *epaR* regarding daptomycin susceptibility.

**Figure 4.**
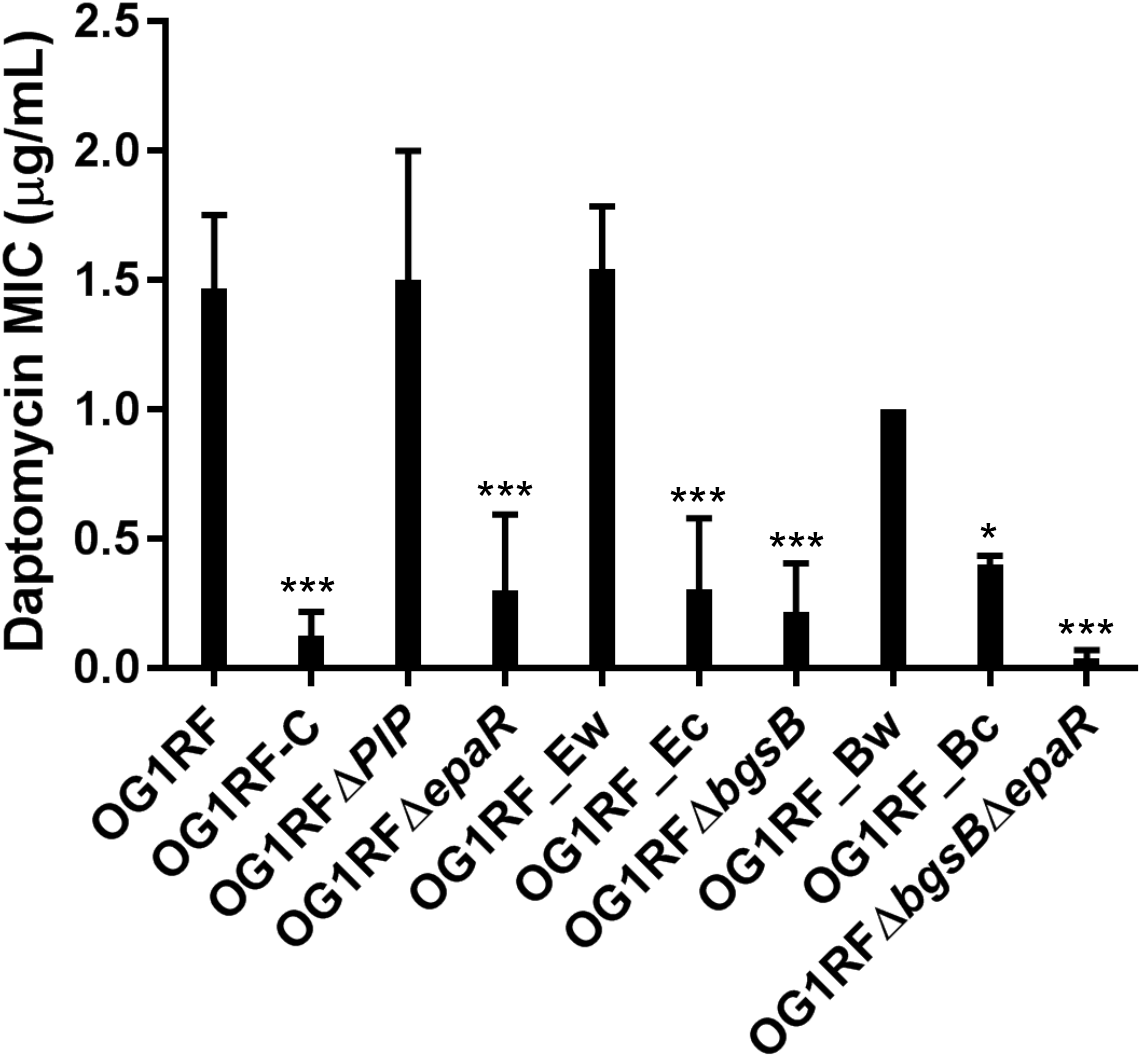
Daptomycin MICs of *E. faecalis* OG1RF and derivatives. Daptomycin MIC was determined by Etest. Data are the average of at least three independent trials. For statistical analysis, DAP MICs were compared to that of wild-type OG1RF. ***, p <0.001; and *, p< 0.05. Note that the names of complemented strains (see Table 1) have been shortened for figure clarity

### *epaR* mutants have increased sodium chloride stress susceptibility

The *epa* gene cluster was up-regulated when *E. faecalis* V583 was grown with 6.5% sodium chloride supplementation, indicating that the Epa polymer has a role in osmotic stress response (25). As such, we investigated the effect of sodium chloride on our *epaR* mutants. We tested our mutants for their tolerance for sodium chloride stress using BHI agar supplemented with sodium chloride at concentrations of 0%, 2.5%, 5%, and 7.5%. Overnight cultures in stationary phase were serially diluted and spotted on these agars. We observed fewer CFU for OG1RF-C, OG1RFΔ*epaR*, and OG1RF *ΔepaRΔbgsB* compared to the wild-type at 7.5% sodium chloride after 72 h incubation (Figure 5). When sodium chloride concentrations of 5% or lower were used, no effect on growth was observed.

**Figure 5.**
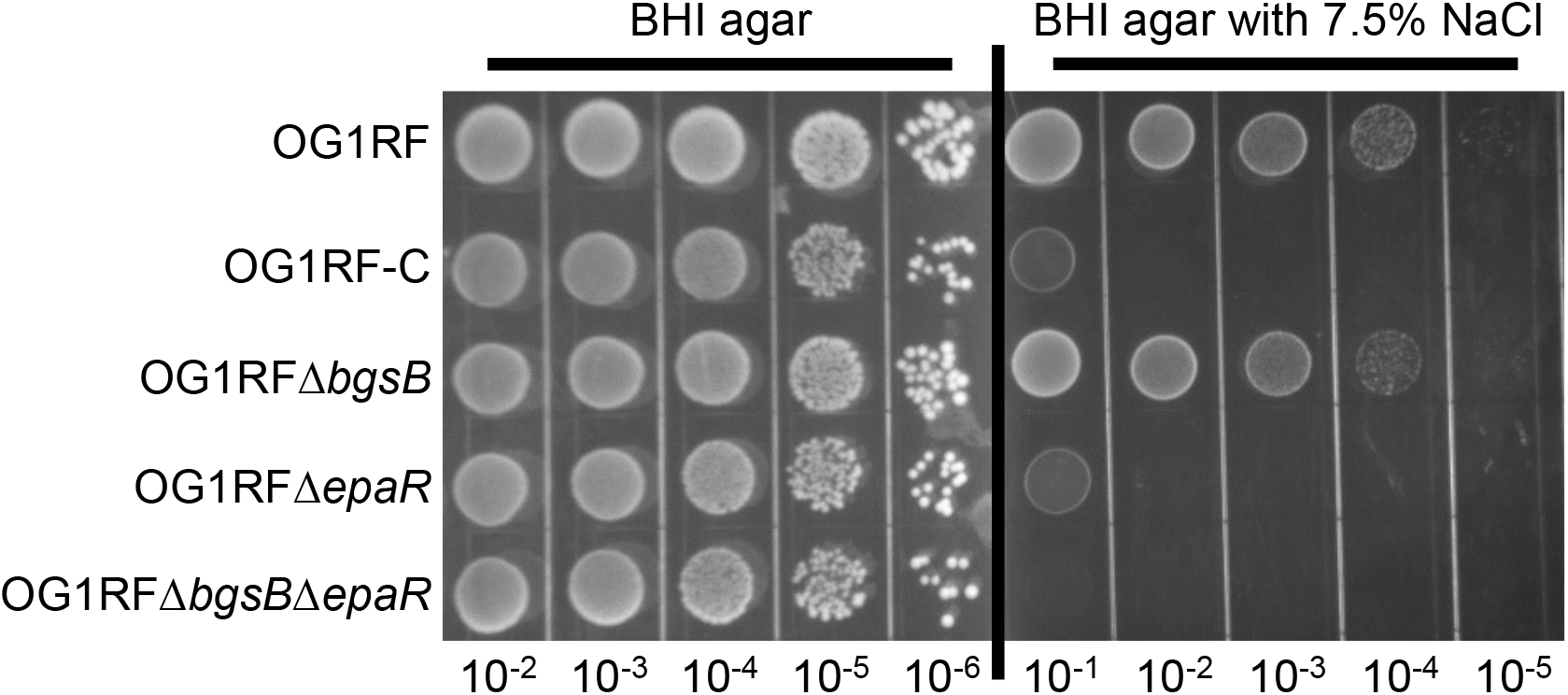
Susceptibility to sodium chloride-induced osmotic stress. Overnight cultures were diluted in PBS and spotted on BHI plates with or without sodium chloride. Images were taken after 72 h incubation. The image shown is representative of three independent trials.

### Multiple different spontaneous *epaR* mutations confer ϕNPV1 resistance

We isolated 9 spontaneous ϕNPV1-resistant mutants of OG1RF and sequenced the *epaR* region of these mutants. All 9 mutants have non-synonymous substitutions in *epaR* (Table 3). Since we know that mutations in *epaR* affect daptomycin susceptibility, we also determined the daptomycin MIC of these ϕNPV1-resistant strains and found that all were more significantly more susceptible to daptomycin than the wild-type (Figure S2).

**Table 3.**
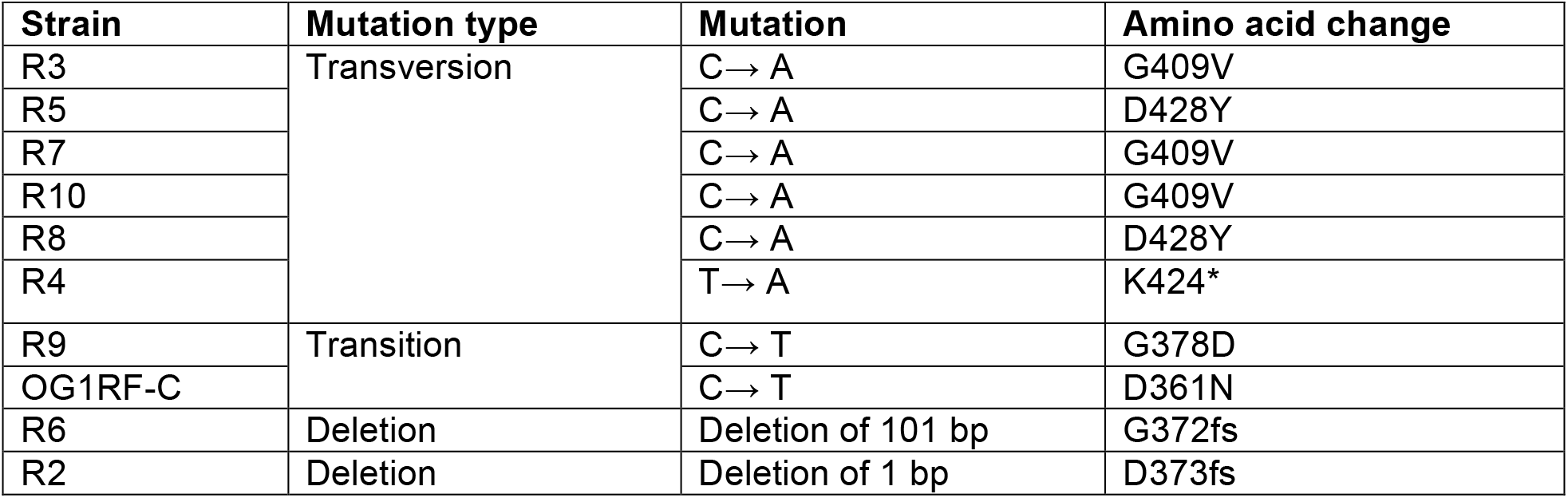
*epaR* variations in spontaneous ϕNPV1-resistant *E. faecalis* OG1RF strains.

| Strain | Mutation type | Mutation | Amino acid change |
| --- | --- | --- | --- |
| R3 | Transversion | C → A | G409V |
| R5 |  | C → A | D428Y |
| R7 |  | C → A | G409V |
| R10 |  | C → A | G409V |
| R8 |  | C → A | D428Y |
| R4 |  | T → A | K424* |
| R9 | Transition | C → T | G378D |
| OG1RF-C |  | C → T | D361N |
| R6 | Deletion | Deletion of 101 bp | G372fs |
| R2 | Deletion | Deletion of 1 bp | D373fs |

Most of the EpaR sequence (amino acid positions 35-458 of 484 total) is a predicted sugar transferase domain (TIGR03025; E-value: 2.45e^−137^). This domain consists of a conserved C-terminal region responsible for the sugar transferase activity (pfam02397) and a variable N-terminal region with predicted flippase activity. All *epaR* mutations in spontaneously ΦNPV1-resistant OG1RF strains occur in the region encoding the sugar transferase domain (Table 3). We identified other proteins containing the same predicted sugar transferase domain as EpaR, and we determined that the altered amino acid positions in our mutants are conserved across most of these proteins (Figure S3).

## Discussion

Due to the high frequency of antibiotic resistance in *E. faecalis*, alternatives to antibiotics, such as phage therapy, are of increasing interest in the United States. In this study, we investigated mechanisms for spontaneous phage resistance in *E. faecalis*. We have reported here that *epaR* is indispensable for ϕNPV1 adsorption to *E. faecalis* OG1RF, and that inactivating mutations in *epaR* constitute a major pathway for ϕNPV1 resistance in this strain background. We also found that inactivating mutations in *epaR* and *bgsB* result in increased susceptibilities to daptomycin and sodium chloride stress. Our results show that resistance to ϕNPV1 comes at a cost.

Adsorption to the host is the first step to a productive phage infection. For a tailed phage particle to successfully adsorb to the host, the tail apparatus on the phage must recognize the corresponding receptor(s) on the host cell surface. When challenged with a high phage titer in a resource-limited environment, receptor mutations in host cells are favored over use of intracellular defense mechanisms (26). This preference for receptor mutations is an especially important consideration in the design of phage cocktails, as using phage that recognize the same receptors could result in decreased efficacy of the treatment (18). Receptors for enterococcal phages have not been well studied. We and collaborators recently identified PIP as a receptor for phage ϕVPE25 and ϕVFW, but PIP is not the sole player in host cell recognition, as ϕVFW and ϕVPE25 can still adsorb to a PIP deletion strain (15). PIP may act as a DNA channel as is implicated in studies of *L. lactis* (14).

The *epa* gene cluster is involved in the synthesis of a cell wall rhamnose polysaccharide referred to as Epa. There is precedence for cell wall rhamnose polysaccharides as phage receptors. The structure of the rhamnose polysaccharide dictates phage host range in *L. lactis* and S. *mutans* (27, 28). In *E. faecalis*, the *epa* gene cluster consists of 18 core genes (*epaA* to *epaR*) and a set of strain-variable genes that occur downstream (20, 29). Unfortunately, the Epa structure has not been determined (30), which is a critical gap in knowledge about the enterococcal cell surface.

The connection between the *epa* gene cluster and ϕNPV1 resistance was first investigated by Teng et al, who assessed ΦNPV1 susceptibilities of *E. faecalis* OG1RF mutants with disruptions in *epaA, epaB, epaE, epaM*, and *epaN* (20). No ϕNPV1 plaques were obtained for *epaB, epaE, epaM*, and *epaN* mutants, and plaque production was reduced by 50% in the *epaA* mutant as compared to the wild-type. However, when the wild-type strain and the *epaA* and *epaB* mutants were assessed for ϕNPV1 adsorption, no differences were noted. Teng et al also examined the polysaccharide content of their mutants and found that production of the ‘P1’ product was absent in the *epaB, epaM, epaN* and *epaE* mutants, but a new polysaccharide product referred to as ‘PS12’ was synthesized. For the *epaA* mutant, both P1 and P12 were produced. The results from Teng et al. suggest that a complete Epa product is required for productive ϕNPV1 infection. Our results support this conclusion as ϕNPV1 does not absorb to our *epaR* mutants, nor do *epaR* mutants synthesize the P1 (or P12) product. However, the P1 product may not be the only requirement for ϕNPV1 adsorption because no significant decrease in PFU was observed when ϕNPV1 was pre-incubated with crude polysaccharide extracts from OG1RF strains with either wild-type or mutant *epaR* (Figure S1). Alternatively, the availability of P1 to the phage may differ in whole cells versus crude extracts.

We investigated the daptomycin susceptibilities of our NPV1-resistant strains with *epaR* mutations because a mutation elsewhere in the *epa* locus was previously linked to daptomycin susceptibility in *E. faecalis*. Specifically, Dale et al. reported that deletion of *epaO* results in increased daptomycin susceptibility (23). Daptomycin is a lipopeptide antibiotic that is used to treat certain Gram-positive bacterial infections (31). The mechanism of action for daptomycin in *B. subtilis* begins with daptomycin binding to the cell membrane, and ultimately leads to the displacement of membrane-associated proteins essential for cell wall and phospholipid biosynthesis (32). In our study, we found that inactivating mutations in *epaR* lead to increased daptomycin susceptibility in *E. faecalis*. The loss of the Epa polymer results in defects in cell wall architecture (20, 33), suggesting that this polymer plays a critical role in enterococcal cell surface physiology. More research on the Epa polymer is required to mechanistically assess its contribution to antibiotic susceptibility in enterococci. In our study, deletion of *bgsB* also resulted in increased daptomycin susceptibility in *E. faecalis*. Deletion of *bgsB* results in loss of glycolipids in the membrane, a longer chain length in the LTA, and increased charge density of the membrane (21). A higher charge density might contribute to daptomycin susceptibility through charge-charge interaction with the calcium-bound daptomycin, but this is speculative. Note that a weakness of our study is that we did not evaluate whether susceptibilities to other antibiotics are altered concomitantly with spontaneous ϕNPV1 resistance or as a result of *epaR* or *bgsB* deletion. Therefore, we cannot comment on whether the altered antibiotic susceptibilities of these strains are specific to daptomycin or are a general defect potentially related to altered membrane/cell wall permeability.

In summary, in this study we characterized a mechanism for spontaneous ϕNPV1 resistance in *E. faecalis* OG1RF and demonstrated that *in vitro* spontaneous ϕNPV1 resistance is accompanied by fitness trade-offs including altered susceptibilities to an antibiotic and to osmotic stress. Experiments for future work include determining the host range of ϕNPV1, and whether other enterococcal phage use the Epa polymer as a receptor for *E. faecalis* adsorption. A critical experiment in terms of possible therapeutic application of ϕNPV1 is to determine whether ϕNPV1 resistance arises *in vivo* (i.e. in the gastrointestinal tract, or during experimental treatment of an *E. faecalis* infection using phage therapy) by the same mechanism as *in vitro*. If ϕNPV1 resistance arises *in vivo* by *epaR* or other *epa* locus inactivation, and this confers an *in vivo* fitness cost to *E. faecalis*, then ϕNPV1 and/or other Epa-targeting phage could be of utility for anti-*E. faecalis* therapies.

## Material and Methods

### Bacterial strains, media, and bacteriophages

A complete list of bacterial strains and bacteriophage used in this study can be found in Table 1. *E. faecalis* strains were cultured in brain heart infusion (BHI) at 37°C without agitation. *E. coli* strains were cultured in LB broth at 37°C with shaking at 225 rpm unless otherwise stated. Plates of the appropriate media were made by adding 1.5% agar to the broth prior to autoclaving. For MIC testing, Muller Hinton media supplemented with 1.5% agar (MHA) was used. Phages were stored in phage buffer as previously described (34). Chloramphenicol (Cm) was used at a concentration of 15 μg/mL when required for selection. 5-bromo 4-chloro-3-indolyl-β-D-galactopyranoside (X-Gal) was used at 120 μg/mL and 40 μg/ml for *E. faecalis* and *E. coli*, respectively. The upper soft agar for phage assays was M17 medium supplemented with 0.75% agar, while the lower layer was BHI supplemented with 1.5% agar.

### Routine molecular techniques and DNA sequencing

Routine PCR reactions were performed using Taq polymerase (NEB) per the manufacturer’s instructions. Phusion polymerase (Fisher) was used for cloning procedures per the manufacturer’s instructions. Plasmid was purified using the GeneJET Plasmid miniprep kit (Fisher). Genomic DNA was isolated using the Ultraclean microbial DNA isolation kit (MoBio). Restriction enzymes, Klenow fragment, T4 polynucleotide kinase (PNK), T4 DNA ligase, and calf intestinal phosphatase (CIP) from NEB were used as instructed by the manufacturer. DNA sequencing was performed at the Massachusetts General Hospital DNA sequencing facility. A complete list of primers used in this study can be found in Table S2.

### Growth curves

Growth curves were performed in triplicate with a Biotek Synergy microplate reader essentially as previously described (35). Overnight broth cultures were diluted 1:1000 into fresh BHI broth and aliquoted into 96 well plates. The optical density at 600 nm of cultures was monitored for 20 h.

### Phage spot assay

0.5 mL of an exponentially growing culture was added to 3 mL soft agar and poured onto BHI agar. 10 μL of the of phage mixture to be titered was spotted onto the soft agar. Plaques were counted after 16 h incubation at 37°C, unless otherwise stated.

### Phage propagation and storage

Phage stocks were prepared by mixing 450 μL of an overnight culture of *E. faecalis* OG1RF with ϕNPV1 at an M.O.I of 10^−2^. The mixture was incubated at 37°C for 15 min and subsequently added to 3 mL M17 soft agar maintained at 55°C. The soft agar was then poured onto BHI agar and incubated at 37°C for 18 h. 5 mL of phage buffer was added to the confluently lysed plate and incubated for 20 min at 37°C with shaking at 75 rpm. The lysate was then collected and centrifuged at 16.6 x g for 1 min to remove cellular debris. The supernatant was filtered with a Whatman 0.2 μm filter to obtain the phage stock. The phage stock was stored at 4°C in the dark. Phage titer was determined using phage spot assays.

### Generation of OG1RF deletion mutants

Gene deletion was carried out via the markerless deletion procedure described by Thurlow et al. (36) with some modifications. Briefly, two 1.0 kb regions flanking *epaR* were amplified with primers 1-4 from Table S2. The two amplified products were ligated with an overlap PCR extension through a 21 bp complementary region underlined in Table S2. The approximately 2.0 kb product was purified and digested with BamHI and EcoRI. The digested product was ligated to plasmid pLT06 through restriction sites added on the primers (highlighted in red in Table S2). The ligation product was then purified and electroporated and propagated in *E. coli* EC1000. OG1RF was made electrocompetent using the glycine method (37) (3% glycine) and transformed with 1 μg of the plasmid. OG1RF transformants were screened for successful transformation and subsequently inoculated in BHI supplemented with Cm at 30°C. The culture was diluted 1:100 in BHI and incubated at 30°C for 2 h followed by 42°C for 4 h. Dilutions of the culture were plated on BHI agar supplemented with Cm and X-gal, and large blue colonies were screened for plasmid integration using primers 5 and 19. The positive colonies were then restruck, incubated at 42°C, and screened once again for plasmid integration. Positive clones were cultured in BHI broth at 30°C for 18 h. To counter-select against clones harboring the plasmid, dilutions of the culture were made on MM9YEG agar, and the deletion of *epaR* was determined by colony PCR with primers 5 and 6 after 36 h incubation at 37°C. Clones positive for the deletion were then restruck on BHI agar and screened again using the same primers. Positive clones were verified for plasmid loss by streaking on BHI agar supplemented with Cm. The *epaR* region was sequenced to confirm the deletion. Deletion of *bgsB* was obtained in a similar fashion.

### Complementation

Complementation of *epaR* in an OG1RFΔ*epaR* background was obtained using a similar strategy to deletion. The insert containing the *epaR* gene and flanking 500 bp upstream and downstream regions was amplified from either OG1RF or OG1RF-C using primers 7 and 8. pLT06 was digested with SphI and blunt-ended with Klenow fragment; the blunt-end product was then treated with CIP. The insert was phosphorylated with T4 PNK and blunt-end ligated to pLT06. The plasmid was purified and transformed into EC1000. Clones with the correct insert size were screened, and their plasmids isolated. Subsequent steps for transformation of OG1RF, integration of the plasmid and counterselection on MM9YEG were as described above for the deletion process. Positive clones for the complementation were confirmed with primers 5 and 6 after counter-selection on MM9YEG. The complemented *epaR* allele was verified through Sanger sequencing.

### Assessment of phage resistance

For assessment of ϕNPV1 resistance, 500 μL of an 8×10^9^ PFU ϕNPV1 stock was added to 3 mL M17 soft agar. The mixture was then poured onto BHI agar. Bacterial culture dilutions were spotted on the soft agar and incubated at 37°C for 18 h. ϕNPV1-resistant bacteria grow on ϕNPV1-containing agar. ϕNPV1-susceptible bacteria do not grow on ϕNPV1-containing agar.

### Assessment of sodium chloride stress tolerance

For assessment of osmotic stress tolerance, BHI plates were supplemented with NaCl (0%, 2.5%, 5%, and 7.5%). Overnight cultures of bacteria are serially diluted and spotted on NaCl-supplemented plates. Plates were imaged after 72 h incubation.

### Phage adsorption assay

An overnight bacterial culture was diluted 1:5 in fresh BHI broth. The culture was then equilibrated at 37°C for 20 min in a water bath. ϕNPV1 was added at an M.O.I of 10^−3^. After 15 min, a 1 mL aliquot was centrifuged at 16.6 x g for 1 min at room temperature. 500 μL of the supernatant was collected, and its titer is determined with the phage spot assay. A medium with only phage added (no bacteria) was used as control. Percent adsorption was determined as follow:

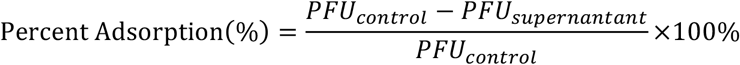

### Isolation of ϕNPV1 resistant mutants

For isolation of a ϕNPV-1 resistant strain from an OG1RFΔPIP background, ϕNPV-1 was used to infect OG1RFΔPIP in a soft agar overlay. The confluently lysed plates were incubated until presumptive phage-resistant colonies arising in the soft agar were observed. These colonies were cultured in BHI broth and used as hosts for ϕNPV1 infection to confirm phage resistance. A confirmed ϕNPV-1 resistant strain, referred to in our study as OG1RF-C, was stocked and used for genome sequencing.

For isolation of ϕNPV-1 resistant strains from an OG1RF background, 500 μL of an overnight culture of OG1RF was infected with ϕNPV1 at an M.O.I of 10^−1^ in a soft agar overlay. 10 colonies that arose on the confluently lysed plate were struck on BHI plates and incubated at 37°C for 18 h. Single colonies from each of the plates were tested for phage resistance by cross-streaking against ϕNPV1. 9 colonies that showed little to no lysis were stocked and used for daptomycin susceptibility testing and *epaR* sequencing.

### Polysaccharide analysis

Polysaccharide extraction was performed as described by Teng et al. with some modifications (20). 200 mL of an overnight culture was centrifuged and resuspended in 750 μL 50 mM Tris buffer, pH 7.5. Mutanolysin (0.25 U/μL) and lysozyme (5 mg/mL) were added to the suspension. The suspension was incubated at 37°C for 2 h. Subsequently, 10 mM MgSO_4_, 2.5 mM CaCl_2_, 0.15 mg/mL DNase I, and 0.15 mg/mL RNase A were added. After an additional 2 h incubation at 37°C, the suspension was centrifuged, and the cellar debris discarded. Proteinase K (100 μg/mL) was added to the clear supernatant, and the mixture was incubated for 16 h. Afterwards, the supernatant was extracted twice with chloroform-phenol-isoamyl alcohol (Sigma Aldrich) and once with chloroform. Ethanol was added to a final concentration of 80% to precipitate the polysaccharide. The precipitate was collected by centrifugation and air-dried. The pellet was resuspended in 50% acetic acid (v/v) in deionized water, and the insoluble material was removed by centrifugation. 30 μL was loaded onto an 1% agarose gel and electrophoresed for 30 min at 130 V. The gel was soaked in staining solution containing Stains-All (Alfa Aesar) and left overnight with gentle rocking. The staining solution was 25% isopropanol, 10% formamide, 65% water, and 0.005% Stains-All. After 18 h, the gel was destained under light for 40 min prior to visualization.

### Daptomycin MIC

Daptomycin MIC was assessed using Etest strips (BioMérieux). 3-5 colonies of similar sizes were resuspended in 500 μL BHI broth and distributed evenly over a MHA plate using a sterile cotton swab. Please note that this inoculation method deviates from clinical susceptibility testing criteria in that we did not determine the CFU of our inocula nor normalize the density of the inocula to a McFarland standard. A daptomycin Etest strip was placed onto the plate, and the plate was incubated for 18 h at 37°C. MIC was determined by recording the number closest to the zone of inhibition. The MIC reported for each strain is the average of at least three independent trials. For trials in which the daptomycin MIC was below the detection limit of the strip (<0.016 μg/mL), the MIC was reported as 0.008 μg/mL for the purposes of statistical analysis. Data were analyzed using the two-tailed unpaired Student’s t test.

### Whole genome sequencing and analysis of OG1RF-C

OG1RF-C genomic DNA was isolated from overnight broth culture using the Ultraclean Microbial DNA Isolation kit (Mo Bio) per the manufacturer’s instruction. The gDNA was sequenced using MiSeq with 2 x 150 bp chemistry at MR DNA (Shallowater, Texas). After sequencing, the reads were mapped to the complete OG1RF reference (NC_017316.1) using CLC Genomics Workbench (Qiagen). Putative mutations were detected using the basic variant detector in CLC Genomic Workbench. Variants occurring at ≥50% frequency in the read assembly and resulting in non-synonymous substitutions were confirmed with Sanger sequencing. BlastP and NCBI Conserved Domains were used to analyze conserved domains in proteins. Amino acid alignment was performed with CLUSTALW (38). Transmembrane helices were predicted with TMHMM version 2.0 (39).

### Accession numbers

The Illumina reads for OG1RF-C have been deposited in the Sequence Read Archive under accession number PRJNA450206.

## Acknowledgments

This work was supported by Public Health Service grant R01AI116610 to K.L.P. Khang Ho was supported by a University of Texas at Dallas Undergraduate Research Fellowship. We gratefully thank Dr. Gary Dunny for providing phage NPV1.

---

**Figure S1:**
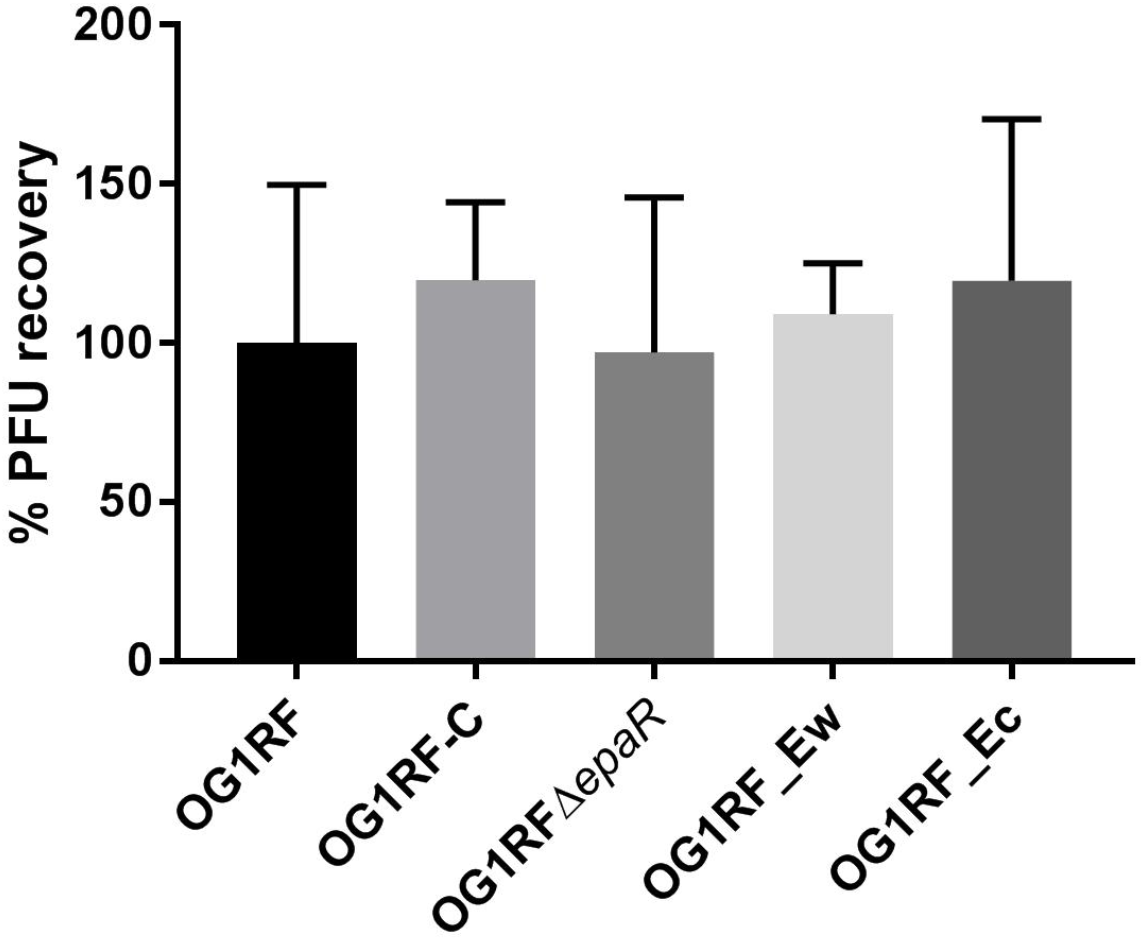
Percent PFU recovery from pre-incubation of phiNPV1 with polysaccharide extract. Polysaccharide extract was prepared as described above except for the final solvent being phage buffer rather than 50% acetic acid. phiNPV1 was added at a PFU of 10^5^ and the mixture was incubated for 15 minutes at 37°C. The PFU was evaluated with phage spot assay. The percent PFU recovery was calculated using a phage buffer only as the base value. The experiment was performed in duplicate.

**Figure S2:**
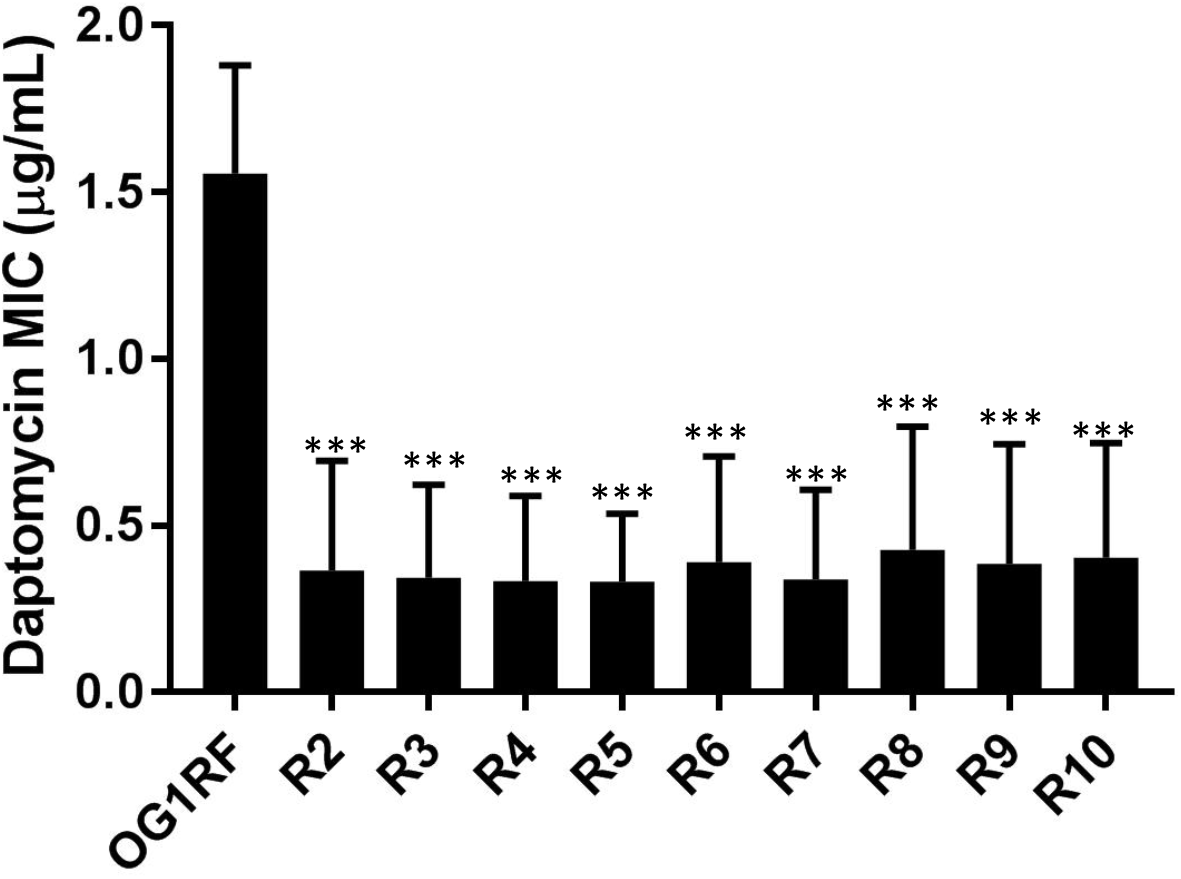
Daptomycin MIC of 9 phiNPV1 resistant strains derived from a confluently lysed plate. 3-5 colonies of indicated strains were resuspended in BHI and swabbed onto a MHB plate. Daptomycin strip was applied to the surface of each plate. The plates were incubated for 18 hours prior to MIC determination. The data represent the average of 6 trials. MIC data points below the detection limit of 0.016 μg/mL were taken to be 0.008 μg/mL for statistical treatment. ***, p <0.001.

**Figure S3:**
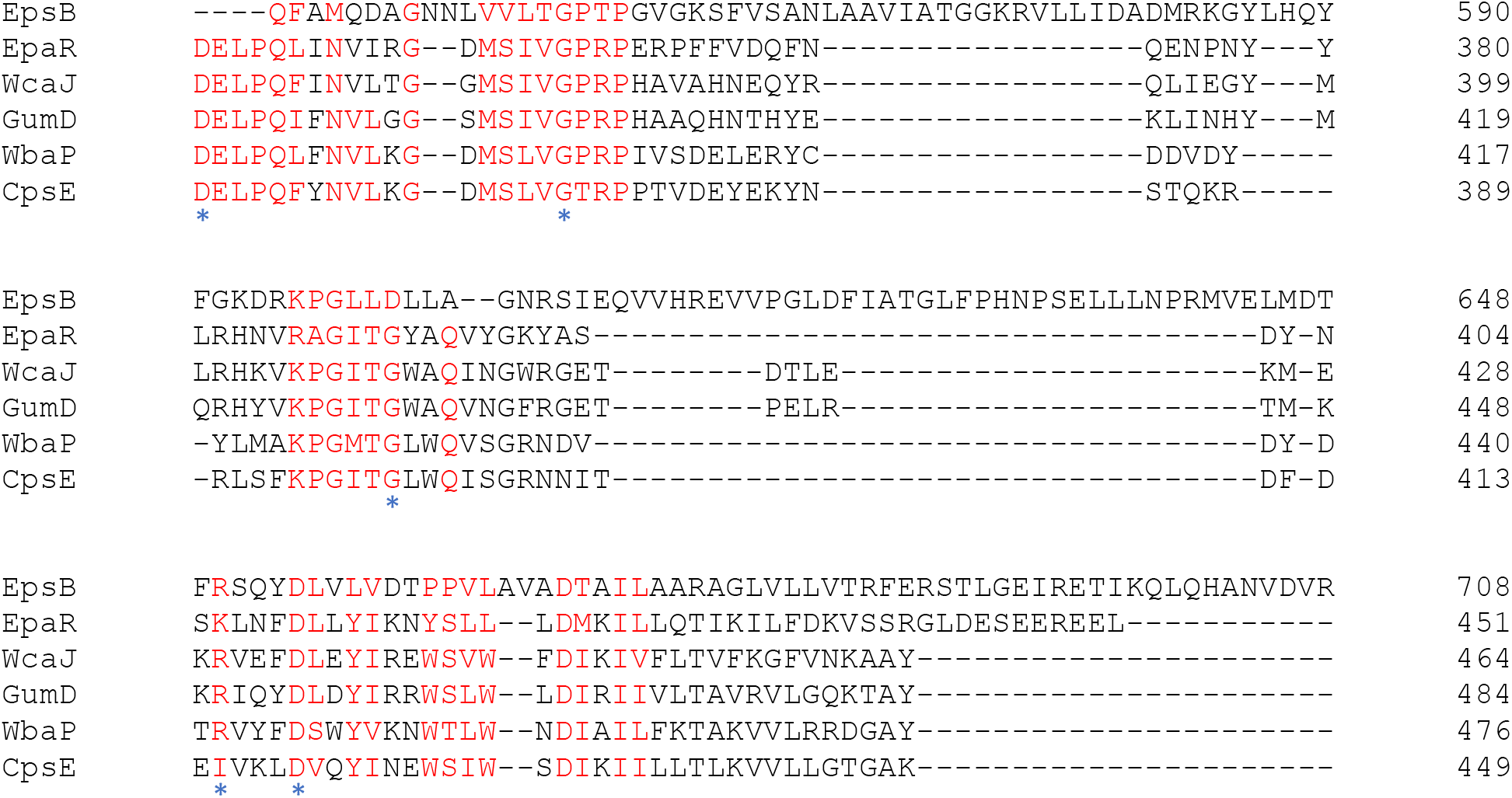
Clustal W alignment of EpaR and its homologues. Conserved residues are highlighted in red. Blue asterisks indicate Blue asterisks indicate where substitutions occurred in phiNPV1 resistant strains of OG1RF.

**Table S1.** Average generation times of *E. faecalis* strains.

| ***E. faecalis* strain** | **Average generation time (minutes)** |
| --- | --- |
| OG1RF | 30.4 |
| OG1RF-C | 33.9 |
| $\Delta epaR$ | 37.4 |
| $\Delta epaR_{Ew}$ | 34.1 |
| $\Delta bgsB$ | 27.8 |
| $\Delta bgsB_{Bw}$ | 39.7 |
| $\Delta epaR \Delta bgsB$ | 42.7 |

**Table S2.**
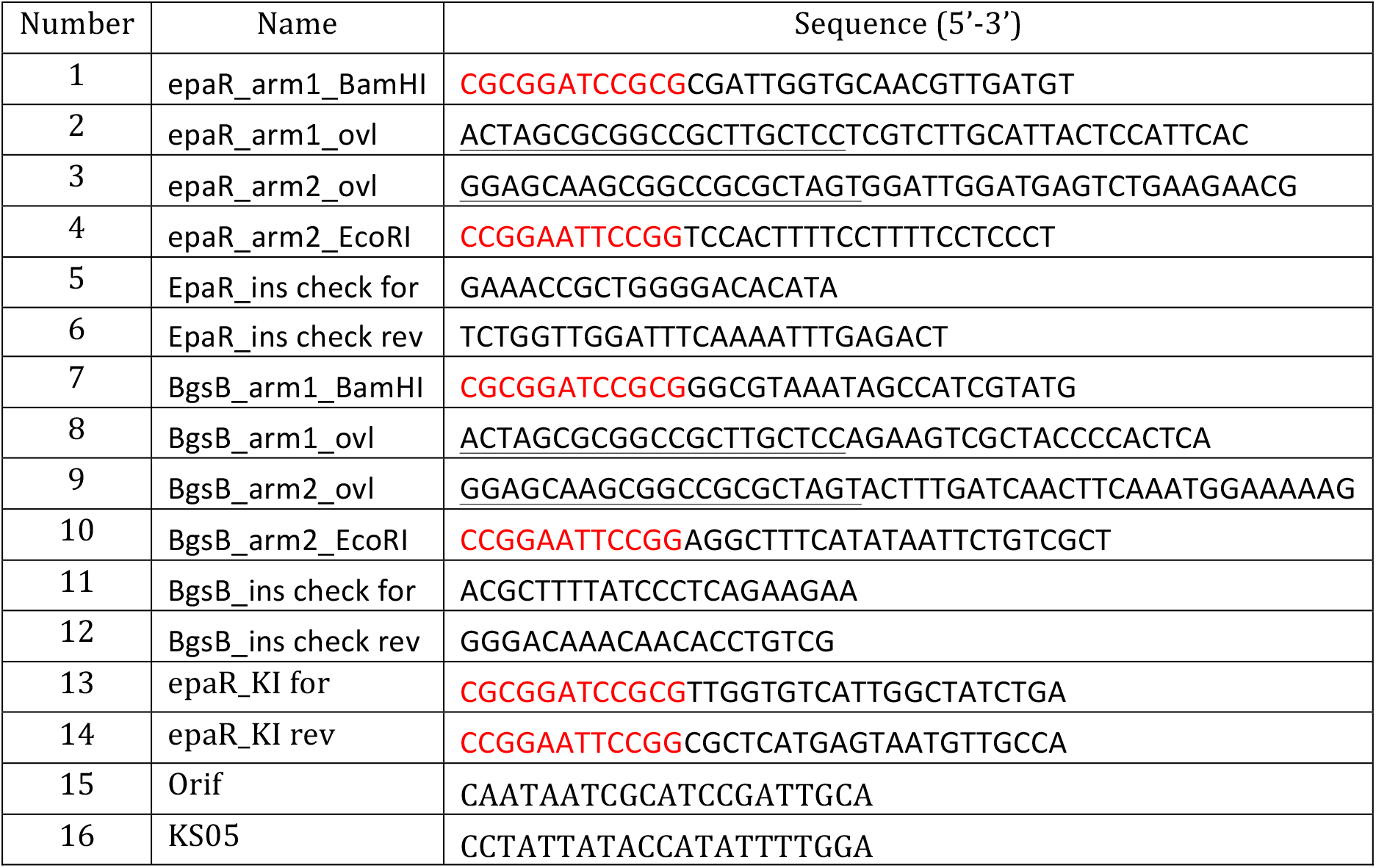
Primers used in this study.

